# Cortical alpha rhythms predictively track occluded motion trajectories

**DOI:** 10.64898/2026.08.21.746169

**Authors:** Susan Ajith, Feyzanur Polat Kamis, Katharina Dobs, Lu-Chun Yeh, Daniel Kaiser

## Abstract

Objects in the real world frequently move along complex, non-linear trajectories shaped by their environment. As they do, they often pass temporarily out of sight, occluded by the surrounding environment. Yet humans can maintain representations of object position and motion even when objects are no longer visible. Here, we investigate the neural basis of tracking objects undergoing non-linear, environmentally constrained motion during occlusion. We recorded EEG while participants mentally tracked the position of a ball falling through differently curved pipes (Pipe condition). The pipe condition was compared to conditions in which only the ball trajectory was visible (Ball-only condition) or in which the pipe was completely covered by an occluder (Occluder condition). On occasional probe trials, the video was terminated at randomized time points and participants had to indicate the ball’s last position using the mouse cursor. Behaviorally, participants systematically reported the ball ahead of its true position, with the strongest forward displacement in the occluded conditions. Neurally, multivariate decoding analysis on the EEG data revealed that alpha-band activity carried direction-specific motion information within and across conditions. Critically, cross-condition decoding revealed shared representations between perceived (Ball-only) and mentally tracked (Occluder) motion with representations in the occluder condition anticipating later representations in the ball-only condition. Together, our behavioral and neural findings provide convergent evidence that during non-linear motion tracking under occlusion, the brain codes object motion ahead of its current position, an ‘ahead of time’ strategy that may underlie temporal precision when tracking objects in a dynamically changing world.

---

The world around us is constantly in motion. Objects move dynamically from one place to another, either by self-propulsion or by physical forces. We often know that objects move even when we cannot directly see them due to (partial) occlusion. During occlusion, humans maintain a representation of the occluded objects’ existence and motion, an ability known as object permanence.

Object permanence is found to rely on maintaining as well as updating the representation of a moving object (Bosco et al., 2015; Shuwairi et al., 2007). Consistent with this, recent studies have shown robust neural evidence for a maintenance and updating mechanism by successfully decoding object position and direction during occlusion (Teichmann et al., 2022; Zbären et al., 2023; Yeh et al., 2024). However, previous work has largely focused on simple one-dimensional or circular motion (Shuwairi et al., 2007; Turner et al., 2025), whereas real-world motion is inherently more complex. Objects often follow non-linear trajectories either because they are self-propelled, or because environmental layouts force them onto constrained paths (e.g., path layouts in natural environments). Recent work suggests the brain can handle both kinds of complexity: it can infer parabolic free-fall trajectories under occlusion (Zbären et al., 2023), and anticipate where linearly moving objects will stop when physical obstacles constrain their path (Yeh et al., 2024). The movement of occluded stimuli is most prominently coded in alpha-band activity (Yeh et al., 2024), consistent with the proposed role of alpha rhythms in the top-down reactivation of visual representations (Xie et al., 2020; Stecher et al., 2025). However, relatively little is known about how the brain tracks motion under more naturalistic constraints - when it is non-linear and context-dependent.

Despite growing evidence for object tracking during occlusion, the neural dynamics governing non-linear, environmentally constrained motion remain poorly understood. EEG studies have largely been restricted to linear or highly predictable motion, and while cross-decoding analyses (i.e., visible to occluded) have demonstrated that tracking during occlusion is supported by neural representations similar to those evoked by direct visual perception, these representations appear largely confined to the early stages of occlusion (Teichmann et al., 2022; Yeh et al., 2024). At later time points, neural activity shifts towards representing the location where the object will reappear (Teichmann et al., 2022). However, this rapid decay may be a hallmark of tracking linear (or highly predictable) motion. In linear trajectories, key motion properties remain stable throughout occlusion, allowing future states to be inferred from the pre-occlusion motion. Non-linear motion, by contrast, involves continuously changing dynamics that render such extrapolation insufficient and require the visual system to update representations during occlusion. Whether and how the visual cortex achieves this for non-linear, environmentally constrained motion therefore remains an important open question.

Here, we investigated object tracking under occlusion when objects move on a non-linear path dictated by the context. Specifically, participants viewed rendered videos of a ball falling through differently curved pipes (pipe condition; see Fig. 1), along with variations where the ball trajectory was shown in isolation (ball-only condition) or where the pipe was hidden by an occluder (occluder condition). In the ball-only condition, the ball was visible, but its trajectory was not known in advance; in the occluded condition, both the ball and pipe were occluded, but the trajectory was indicated by the color of the occluder. Participants were asked to mentally track the position of a falling ball and report its position on occasional target trials. EEG was recorded throughout the experiment, and analyses focused on alpha-band activity given its established role in predictive motion extrapolation (Turner et al., 2023; Yeh et al., 2024). Our behavioral results show that participants systematically reported the ball ahead of its true position, with the strongest effects in the occluded conditions. Our neural results demonstrate robust alpha representations of the ball’s motion direction in each condition. Cross-decoding analyses reveal shared neural representations between visible ball motion without prior trajectory constraints (ball-only condition) and occluded motion with known trajectory constraints (occluded condition). Critically, representations in the occluded condition anticipated later representations in the ball-only condition, revealing anticipatory coding of ball motion during occlusion. Together, our neural and behavioral findings reveal that alpha-band dynamics predictively track non-linear occluded motion when the trajectory is implied by contextual information, providing new insight into how the brain maintains temporal precision in a dynamically changing world.

## Methods

### Participants

Forty-one volunteers (25 female; mean age = 26.27, SD = 4.65) participated in the experiment. All participants had normal or corrected-to-normal vision. They provided informed written consent and received 10 euros per hour for their time. The study was approved by the Ethics Committee of the Justus Liebig University Giessen and was in accordance with the 6th Declaration of Helsinki.

### Stimuli

Stimuli consisted of 6s video clips (960 × 540 pixels; 24.6° × 14.3° visual angle) depicting a ball falling under gravity (g=9.81m/s2) through 4 pipes of different shapes (**Pipe condition**) (see fig 1B). The shapes of the pipe followed four possible diagonal trajectories: up-right, up-left, down-right, or down-left. Then the stimuli were modified in two ways: 1. The pipe was rendered invisible, such that only the ball was visible as it moved through the invisible pipe (**Ball-only condition**), 2. The pipe and ball were completely occluded by an object (**Occluder condition**). In this condition, the color of the occluder indicated the shape of the pipe behind it: Light Green (up-right), Light Red (up-left), Dark Green (down-right), and Dark Red (down-left). All stimuli followed a standardized temporal sequence: the ball initially descended vertically for 2.2s, changed direction, and continued along its assigned diagonal path until 5s.

Stimuli were generated using Blender (v4.4.0) and rendered at a resolution of 1920x1080 pixels (down-sampled to 960 × 540 pixels for presentation). Each 6s clip was presented at 24 fps, subtending a visual angle of 24.6° × 14.3°. The pipes subtended a visual angle of ∼8.9° × ∼2.0°. Leftward trajectories (up-left and down-left) were animated natively, while rightward trajectories (up-right and down-right) were created by horizontally flipping the leftward animations to ensure spatial symmetry. For the Ball-only condition, the pipes in the respective directions were rendered invisible. For the Occluder condition, occluders of assigned colors were superimposed onto the respective pipe videos. This procedure resulted in a total of 12 unique video stimuli (3 conditions × 4 motion directions).

**Fig 1.**
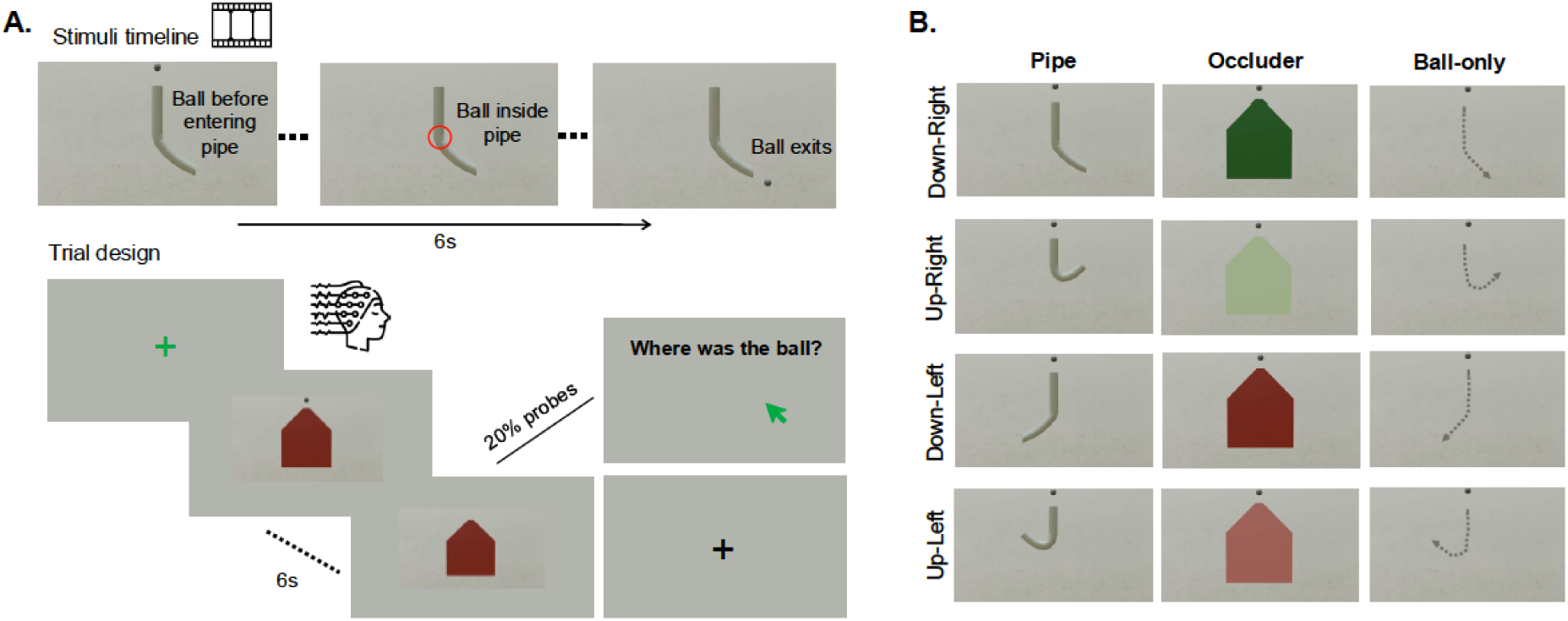
Experimental design. **(A)** Stimuli consisted of 6s video clips depicting a ball falling under gravity through 4 pipes of different shapes (see fig 1B). The shapes of the pipe followed four possible trajectories: up-right, up-left, down-right (i.e., example stimulus), or down-left. All stimuli followed a standardized temporal sequence: the ball initially descended vertically for 2.2s, changed direction, and continued along its assigned diagonal path until 5s. In the main experiment, each trial began with a central fixation cross. Stimulus onset was triggered by participants maintaining fixation. On 20% of trials (probe trials), the video paused at a randomized frame and was replaced by a grey screen; participants then used a mouse cursor to click on the last perceived position of the ball. No overt response was required on the remaining non-probe trials. **(B)** Motion trajectories were presented across three conditions. In the *Pipe* condition, the ball is initially visible until it enters the pipe structure and then reappears after the ball exits the pipe. In the *Occluder* condition, the ball and the pipe is occluded by a centrally positioned opaque occluder. Here, the color of the occluder indicated the shape of the pipe behind it: Light Green (up-right), Light Red (up-left), Dark Green (down-right), and Dark Red (down-left). In the *Ball-only* condition, the pipe was rendered invisible, and only the ball was visible throughout its motion.

### Experimental paradigm

Participants sat 60 cm from the 32-inch screen using a chinrest to maintain a consistent viewing distance and minimize head movement. We instructed participants to mentally track the movement of the ball while maintaining central fixation. EEG signals were recorded during the experiment.

Prior to the experimental blocks, participants were briefed on the association between the colour of the occluder and the corresponding ball trajectory. They were required to memorize these mappings and successfully recall them before proceeding to the practice block. The practice block consisted of one probe and one non-probe trial for each condition and trajectory, resulting in a total of 24 trials. This ensured participants could maintain central fixation, respond accurately to probes, and apply the color-trajectory associations; the practice block was repeated if these criteria were not met.

The main experiment consisted of five blocks of 144 trials each. Within each block, 20% of trials were probe trials. In these trials, the video paused at a randomized frame and was replaced by a grey screen; participants then used a mouse cursor to click on the last perceived position of the ball. In non-probe trials, no overt response was required while the video was being played. Inter-trial intervals (ITI) were fixed at 500 ms, and the total session duration was approximately 60 minutes.

To ensure central fixation and minimize ocular artifacts, the experiment strictly enforced maintaining central fixation during the trials. Each trial started when participants maintained gaze within a central 3.1° × 3.1° box for 200 ms. If a participant’s gaze moved beyond this central box at any point during the trial, a warning was shown (“Eyes not centrally fixated”) along with the location of gaze highlighted in red, and the trial was aborted (Teichmann et al., 2023). Aborted trials were appended to a final block of previously aborted trials that was shown after the five experimental blocks.

### Behavioural data analyses

We analysed participants’ responses on the probe trials using two metrics: error magnitude and error direction. Error magnitude was defined as the Euclidean distance (in pixels) between the ball’s true spatial coordinates (in pixels on the screen) and the participant’s reported coordinates during the response window. Error direction was defined as the directional bias in the response. It was calculated as the angle of the vector (in degrees) connecting the ball’s true spatial coordinates and participant’s reported coordinates during the response window. Data were analysed separately for each condition (pipe, ball-only, occluder), calculating error magnitude and direction for each participant and motion trajectory.

For each participant, error magnitude was averaged across motion directions (up-left, up-right, down-left, down-right) within each condition. We then tested whether differences in error magnitude between condition pairs (pipe–occluder, pipe–ball-only, occluder–ball-only) were significant using paired t-tests. False discovery rate (FDR) correction was applied to control for multiple comparisons. To compare mean error directions across the four motion trajectories within each condition, we employed a non-parametric permutation-based circular ANOVA, as the data did not meet the assumptions of circular normality and concentration required for parametric tests (i.e., Watson-Williams test), evidenced by mean resultant vector lengths falling below the 0.45 threshold. A null distribution was generated by shuffling stimulus labels (i.e., motion trajectory labels) within participants across 10,000 iterations. The p-value was calculated as the proportion of permutations yielding a circular variance equal to or greater than the observed circular variance.

#### EEG data acquisition and preprocessing

Electrophysiological data were recorded using an Easycap system with 64 channels and a Brain Products amplifier with 1000 Hz sampling rate. The electrodes included seven sites in the central line (Fz, FCz, Cz, CPz, Pz, POz, and Oz) and 28 sites over the left and right hemispheres (FP1/FP2, AF3/AF4, AF7/AF8, F1/F2, F3/F4, F5/F6, F7/F8, FC1/FC2, FC3/FC4, FT5/FT6, FT7/FT8, FT9/FT10, C1/C2, C3/C4, C5/C6, T7/T8, CP1/CP2, CP3/CP4, CP5/CP6, TP7/TP8, P1/P2, P3/P4, P5/P6, P7/P8, PO3/PO4, PO7/PO8, PO9/PO10, and O1/O2). AFz served as the ground electrode, and Fz served as a reference electrode. EEG data were preprocessed offline using the Fieldtrip toolbox (Oostenveld et al., 2011) in MATLAB (MathWorks). EEG data were re-referenced using the average of all electrodes and epoched from -1 to 5s relative to stimulus onset. Data were band-stop filtered at 48-52 Hz to remove line noise. Epochs were baseline-corrected from -100 to 0ms. Noisy channels were removed by visual inspection and interpolated using the mean signals of the neighbouring channels.

### EEG time-frequency analysis

Time-frequency analysis was performed using the Fieldtrip toolbox. Continuous Morlet wavelet transformation with a 7-cycle length was used for time-frequency decomposition from -1 to 5 seconds in 50-ms steps. Power values in each frequency band (4 - 7 Hz for the theta band, 8 -12 Hz for the alpha band, 13 - 30 Hz for the beta band, and 31 - 70 Hz for the gamma band) were averaged for the following decoding analysis.

### EEG decoding analysis

To track the neural representation during dynamic motion, we conducted a multivariate classification analysis on time-frequency-resolved EEG data using the CoSMoMVPA toolbox (Oosterhof et al., 2016). Separate linear discriminant analysis (LDA) classifiers were employed for each motion axis: vertical (Up vs. Down) and horizontal (Left vs. Right). Separate classification analyses were performed for all time points from video onset to 5 seconds post-onset.

#### Temporal generalization

To examine whether motion-related activation patterns observed at one time point generalise to other time points, we trained classifiers on data from each frequency band from one time point and tested them on data from all other time points in the epoch. This approach results in temporal generalization matrices in which each cell corresponds to a decoding performance at a unique training and testing time combination (Grootswagers et al., 2017; King & Dehaene, 2014). Accuracy was aggregated across all possible train-test splits. Area Under the Curve (AUC) served as the measure of decoding sensitivity, with a theoretical chance level of 0.50.

#### Within-condition temporal generalization

We first assessed motion representations within each condition. In these analyses, training and testing data stemmed from the same condition, resulting in within-condition temporal generalization matrices. For instance, for each condition, we trained a linear classifier to distinguish leftward (up-left and down-left) from rightward (up-right and down-right) trajectories from alpha-power topographies. Cross-validation folds were generated using random five-fold partitions. The number of leftward and rightward trials was balanced across folds. This procedure was repeated five times, resulting in 25 folds per participant. The resulting matrices were symmetrized by averaging across train-test directions (e.g., Train A/Test B and Train B/Test A) for each participant (van den Hurk & Op de Beeck, 2019). Individual matrices were then averaged to produce group-level generalization matrices.

#### Cross-condition temporal generalization

Next, we assessed how motion representations are shared across different motion conditions. Here, we trained a classifier on data from each time point of one condition (e.g., Pipe) and tested it on data from all possible time points of a different condition (e.g., Occluder), resulting in cross-condition temporal generalization matrices. These matrices were symmetrized by averaging across train-test directions (e.g., Train A/Test B and Train B/Test A) for each participant (van den Hurk & Op de Beeck, 2019). Individual matrices were then averaged to produce group-level cross-condition generalization matrices.

### Statistical Testing

To identify time points at which classification performance was significantly above chance while correcting for multiple comparisons, we used a cluster-based permutation test with threshold-free cluster enhancement (TFCE; Smith & Nichols, 2009), generating the null distribution via 10,000 sign-flip permutations (Maris & Oostenveld, 2007). This test was one-sided, testing for AUC significantly greater than chance (0.5). The same test was used to compare decoding performance between conditions.

All data and code are publicly available: https://doi.org/10.5281/zenodo.21922131

## Results

### Behavioural performance indicates predictive motion tracking under occlusion

**Fig. 2.**
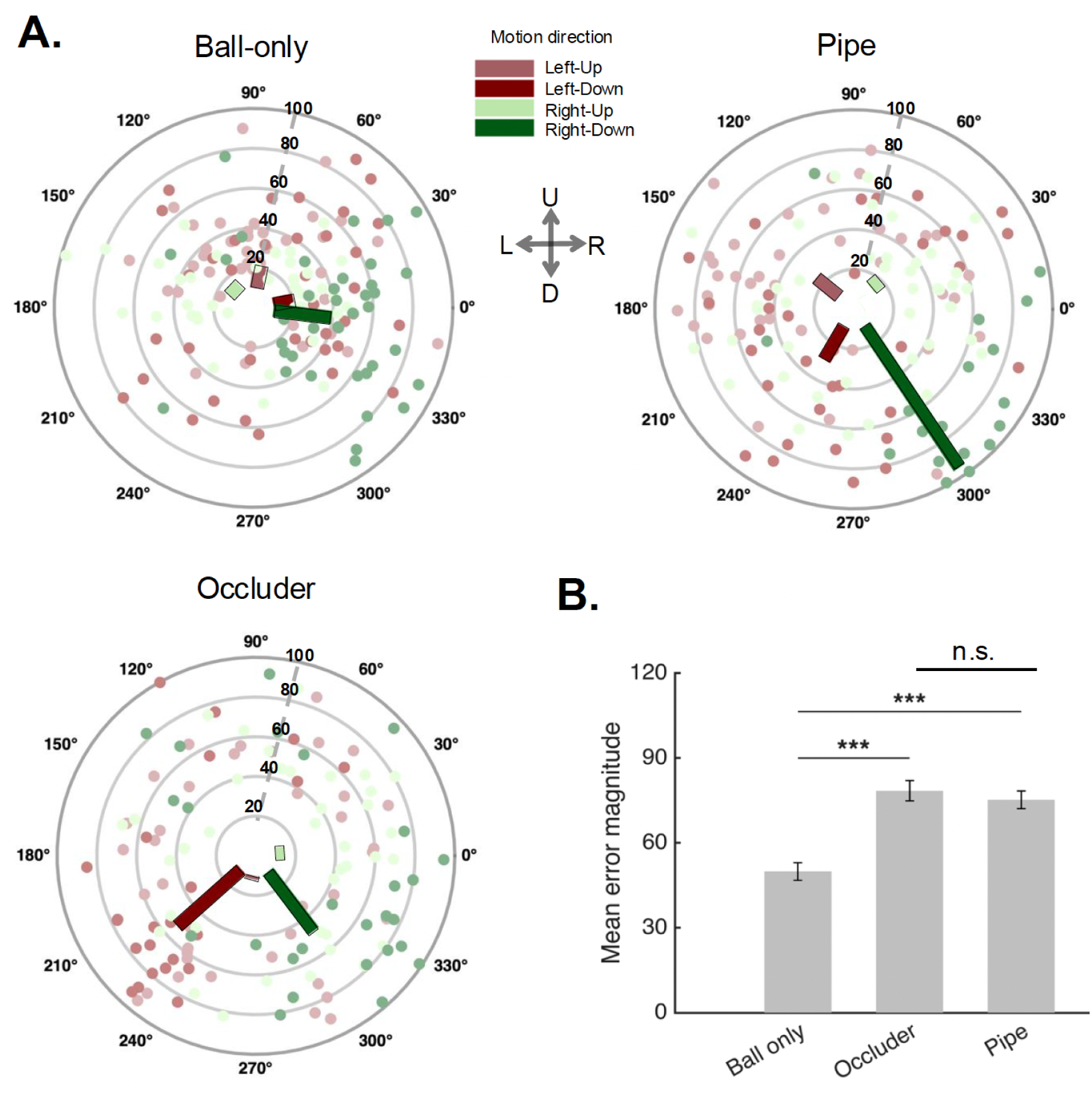
Behavioral performance. On probe trials, participants reported the last position of the ball via mouse click on the screen. Two metrics were assessed: error magnitude (Euclidean distance in pixels between the true and reported ball position) and error direction (angle of the vector between the ball’s true spatial coordinates and participant’s response coordinates). **(A) Error distributions.** Each dot represents a participant’s mean error magnitude and mean error direction for a given motion trajectory. The aggregated vector summarizes the group-level error, where the length reflects the error magnitude and the angle reflects the error direction. **(B) Mean error magnitudes across conditions.** The error magnitude was significantly larger in the Occluder and Pipe conditions than in the Ball-only condition. Asterisks indicate *p* < 0.05. Error bars represent SEM.

On target trials, the video was terminated at randomly chosen time points. Participants subsequently reported the location of the ball at the time when the video was terminated, indicated by a mouse click on the screen. Behavioral responses revealed three effects: First, the mean error magnitude was greater in the pipe and occluder condition, compared to the ball-only condition (both t(40)>11, p<0.001). No significant difference was observed between the occluder and pipe conditions (t(40)=1.9, p=0.05). This indicates greater deviations from the true ball positions when the last ball position cannot be directly perceived but needs to be mentally tracked. Second, we observed a significant difference in error directions across ball trajectories (permutation-based circular ANOVA, p<0.001), with an error direction of -37° for the right-down stimuli, -133° for the left-down stimuli, 30.0° right-up stimuli, and 115° left-up stimuli (mean resultant lengths R = 0.60, 0.16, 0.17, and 0.22, respectively). Third, our results show significantly stronger forward displacement during downward compared to upward motion conditions of the ball (t(40)=10.22, p<0.001). Together, these results indicate that the tracking of occluded motion is more error prone but that errors are not random: they follow the direction of the ongoing motion, revealing a predictive, “ahead of time” representation of the ball’s position during occlusion.

### EEG alpha rhythms carry direction-specific motion information

By performing multivariate decoding analysis on time-frequency-resolved EEG patterns, we tested the extent to which motion information is shared within and across different motion conditions. Our analysis focused on patterns in the alpha power (8–12 Hz) as alpha band activity has been previously associated with motion extrapolation under occlusion (Turner et al., 2023; Yeh et al., 2024). For results with other frequency bands, see Supplementary fig S3.

We trained linear classifiers on alpha activity patterns to discriminate horizontal motion direction (left versus right), separately for the Pipe, Ball-only, and Occluder conditions. Our main analyses focus on the horizontal axis, where stronger effects were expected given the greater distance of the ball positions. Results for the vertical motion axis are shown in the supplementary materials (Supplementary figs S5, S6, S7).

**Fig. 3.**
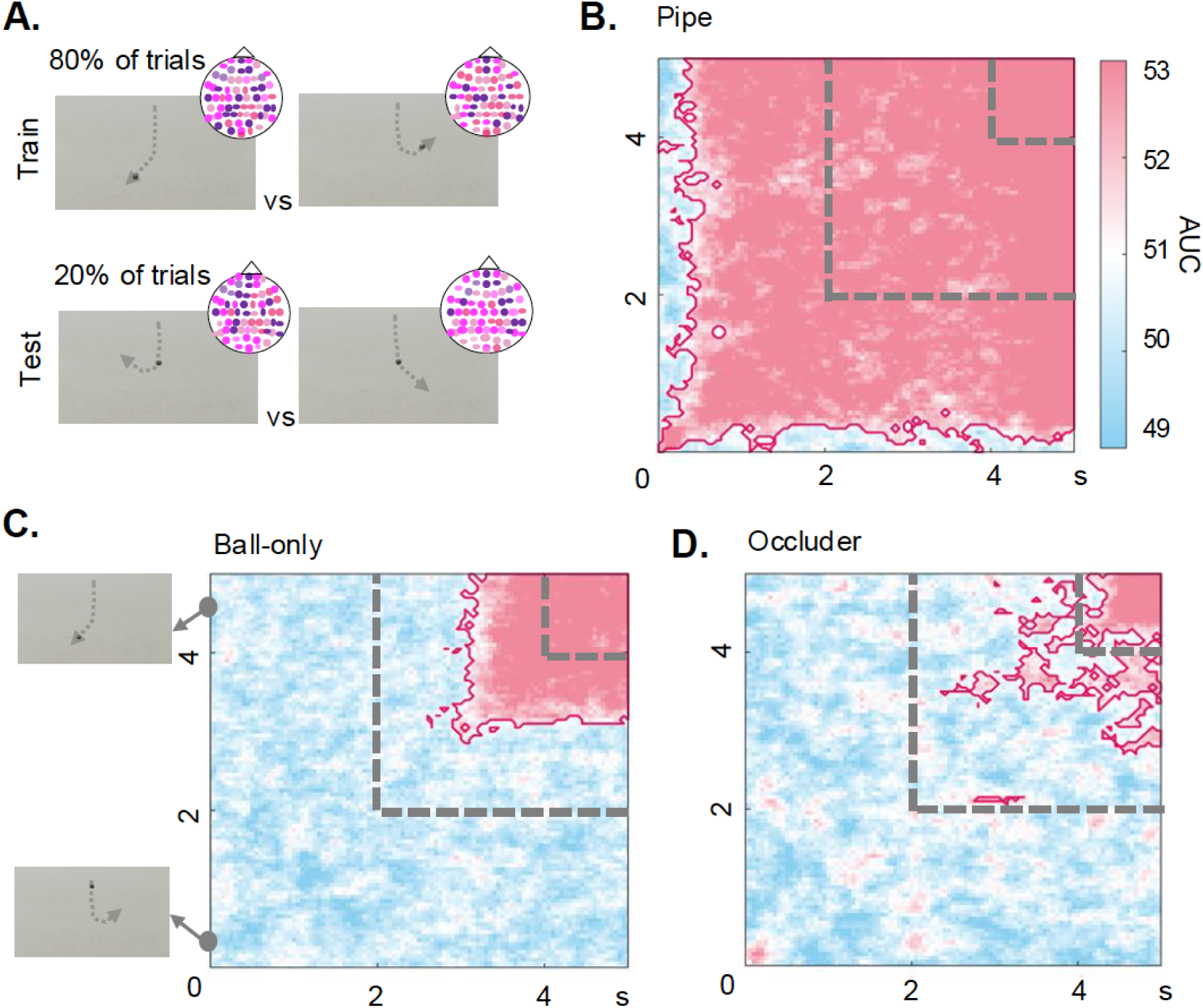
Within-condition temporal generalization. **(A)** Linear classifiers were trained and tested on alpha power (8–12 Hz) patterns to decode horizontal motion direction (left versus right). Here, classifiers were trained and tested within the same conditions. We performed a repeated 5-fold cross validation, where 80% of trials were used for training the classifier and 20% of trials were used for testing. This was repeated 5 times. **(B).** In the pipe condition, direction information remained significant throughout the trial (0-4.5s). **(C)** In the ball-only condition, significant clusters emerged between 2.9s-4.5s, corresponding to the critical time window where direction information became available after the ball departed from its vertical course. Insets on the left show the stimulus frame that participants viewed at the highlighted timepoint. **(D)** In the occluder condition, direction information was significant between 2.4-4.5s suggesting that direction-specific motion representations are observed only once the ball is perceived or inferred to change direction. Each cell in the matrix reflects decoding accuracy (AUC) at a given train–test time point. The grey dashed lines demarcate the critical time window (2–4 s), spanning the period from which the ball changes direction from its vertical descent until the ball exits. Significant clusters (cluster-based permutation test with TFCE, *p* < 0.05) are outlined in dark pink. Note that decoding in the Pipe condition is strongly driven by the presence of the pipe shape throughout the trial.

First, we employed a within-condition temporal generalization approach to map the temporal dynamics of motion direction representations separately for each condition. We were able to successfully decode leftwards versus rightward motion direction from alpha power patterns in all conditions (fig. 3). In the pipe condition, direction information remained significant throughout the trial (0-4.5s) (peak AUC=55% at 3.8s). Note that direction decoding in the pipe condition does not only reflect the ball’s motion direction but mainly the direction of the visible pipe. In the ball-only condition, significant clusters emerged between 2.9s-4.5s, (peak AUC=57% at 3.8s) corresponding to the critical time window where direction information became available after the ball departed its vertical course. In the occluder condition, direction information was significant between 2.4-4.5s, (peak AUC=53% at 3.6s), suggesting that direction-specific motion representations are again observed only once the ball changes direction. Notably, although direction information is available from the outset in this condition (indicated by the colour of the occluder), alpha activity only encodes the motion direction once the ball is perceived or inferred as changing its direction. Comparing this pattern to the early decoding onset in the pipe condition suggests that alpha rhythms more strongly code for spatial information (like the different retinotopic locations of the pipes) than for feature information (like the different colors of the occluders). Furthermore, the shapes of the leftward-versus rightward-bent pipes produce lateralization differences, which alpha rhythms may prominently reflect (Bacigalupo & Luck, 2019). In all conditions, the strong decoding after 4s is related to the ball reappearing from behind the occluder.

### Alpha dynamics carry shared direction-specific information across conditions

To test for shared representations across conditions, we trained linear classifiers to discriminate motion direction (left versus right) using data from one condition and tested them on data from another condition (and vice versa, for all condition combinations; fig 4A).

**Fig 4.**
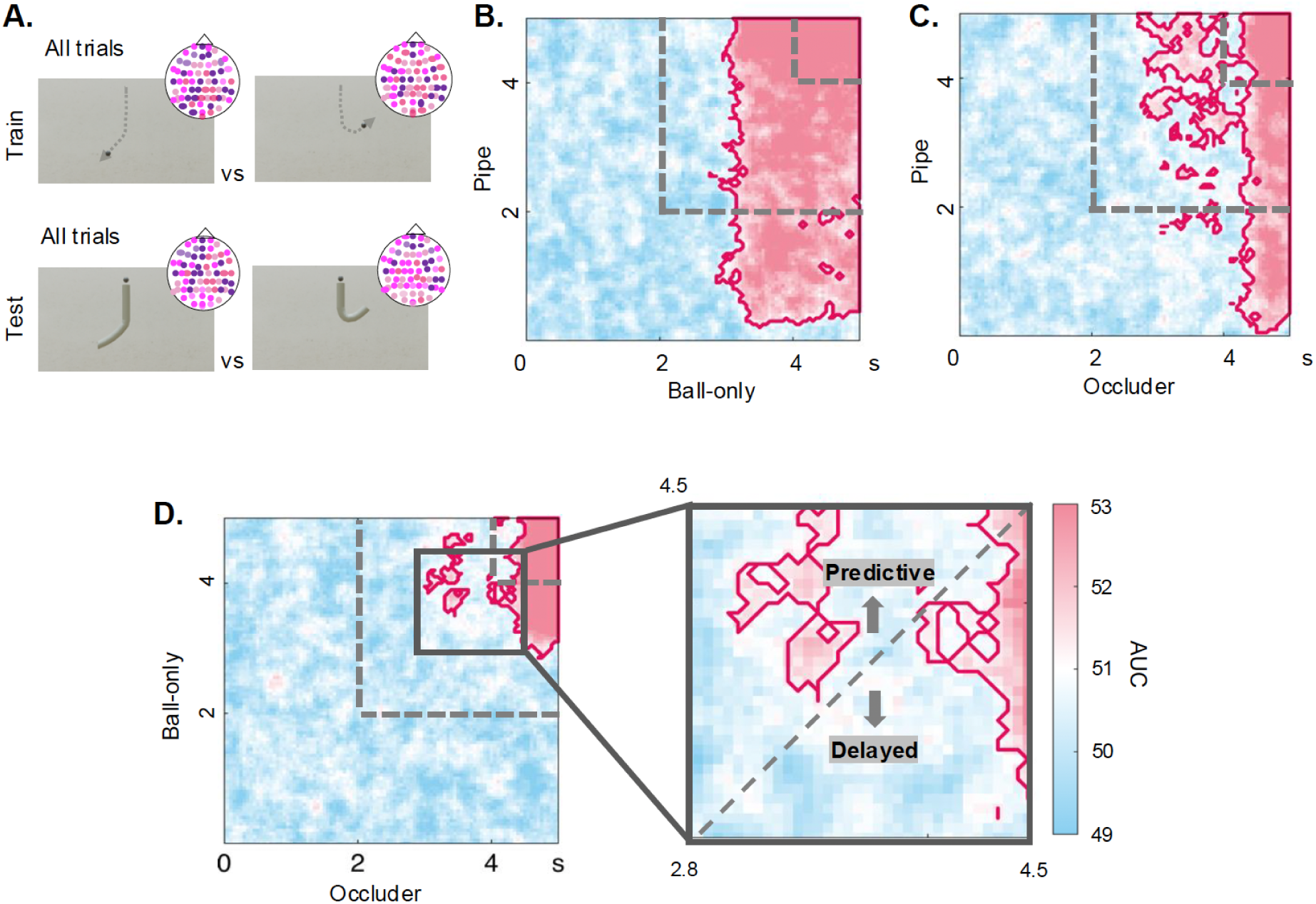
Cross-condition temporal generalization. **(A)** Classifiers were trained on one condition and tested on another (and vice versa, across all condition combinations). **(B)** In the ball-only-to-pipe condition cross decoding, a widespread cluster is observed spanning 0.2s-4s in the ball-only condition and from 2.75s-4.5s in the pipe condition. However, this pairwise combination is confounded by the visual similarity between conditions, as the ball physically overlaps with the pipe once it departs its vertical course. **(C)** In the pipe-to-occluder condition cross decoding, a cluster is observed spanning 1.60s-4.5s in the pipe condition and 2.70s-4s in the occluder condition. This shows that tracking the ball under occlusion follows similar temporal dynamics in both conditions, despite their different constraints: the pipe provides visible path information, whereas the occluder indicates a memorised path. **(D)** In the ball-only-to-occluder condition, significant clusters emerge between 3.5s-4.5s in the occluder condition and 3.5-4.5s of the ball condition. In the inlay, we highlight the temporally specific shared representation prior to ball exit, extending from 3.25s to 4s. The cluster above the diagonal, where occluder training times precede ball-only test times, provides evidence for predictive representation during occlusion. Here, the coding of the ball position in the occluder condition anticipates the future position of the ball in the ball-only condition. Each cell in the matrix reflects decoding accuracy (AUC) at a given train–test time point. Significant clusters (cluster-based permutation test with TFCE, *p* < 0.05) are outlined in dark pink.

First, we trained decoders on data from the pipe condition and tested them on data from the ball-only condition (and vice versa). Here, we observed a widespread cluster spanning 0.2s-4s in the ball-only condition and from 2.75s-4.5s in the pipe condition (peak AUC=54% at 2.9s in the ball condition and 3.8s in the pipe condition). However, this pairwise combination is confounded by the visual similarity between conditions: once the ball departs from its vertical course in the ball-only condition, it physically overlaps with the pipe in the pipe-only condition, strongly driving the direction decoding.

Second, we trained decoders on data from the pipe condition and tested them on data from the occluder condition (and vice versa). Here, we observed a cluster spanning 1.60s-4.5s in the pipe condition and 2.70s-4s in the occluder condition (peak AUC=53% at 3.3s in the pipe condition and 3.9s in the occluder condition). As in both the pipe and occlude conditions, the ball position over time cannot be observed but needs to be inferred from context (either the physical pipe or the color of the occluder), these results show that tracking the ball under occlusion follows similar temporal dynamics in both conditions. Interestingly, this is despite the different constraints imposed by the two conditions, where the pipe provides exact and visible path information that constrains the trajectory, whereas the occluder provides a pointer towards a memorized path.

Third, we trained decoders on data from the ball-only condition and tested them on data from the occluder condition (and vice versa). Here, significant clusters emerged between 3.5s-4.5s in the occluder condition and 3.5-4.5s of the ball condition (peak AUC=52% at 3.8s in the occluder condition and 3.4s in the ball condition). Most interestingly, we observed a temporally specific shared representation prior to ball exit, extending from 3.25s to 4s. This cluster is highlighted in the inlay panel in fig. 4D (right). The cluster primarily spanned time point combinations above the diagonal, with earlier representations in the occluder condition generalizing to later representations in the ball-only condition. This suggests a predictive representation during occlusion, where the coding of the ball position in the occluder condition anticipates the future position of the ball in the ball-only condition. This result is consistent with our behavioral results showing an ahead of time coding when participants are explicitly cued to report the ball position.

## Discussion

In our study, we used EEG to investigate the neural basis of motion tracking under occlusion when objects are forced on non-linear paths through environmental constraints. Participants mentally tracked the position of a ball falling through differently curved pipes (Pipe condition). We further added conditions in which only the ball trajectory but not the pipe was visible (Ball-only condition) or in which the pipe was completely covered by an occluder whose color indicated its shape (Occluder condition). Behavioral data from occasional target trials, during which participants indicated the ball’s current position, revealed evidence for predictive tracking: The ball was placed further along the trajectory than its current position, an effect that was particularly prominent in the Pipe and Occluder condition. Decoding analysis on the EEG data revealed that alpha-band activity carried direction-specific motion information within and across conditions, consistent with the finding that alpha rhythms serve as a neural marker for motion tracking (Stecher et al., 2025; Yeh et al., 2024). Critically, shared representations between the ball-only and occluder conditions revealed predictive neural tracking under occlusion, where earlier time points in the occluder condition represented the ball’s future position in the ball-only condition. Together, our findings demonstrate shared representations across perceived and mentally tracked motion trajectories, even when the trajectories are non-linear but predicted by contextual constraints. We also show that representations during occlusion are predictive, with neural coding operating “ahead of time”.

In our experiment, motion trajectories were constrained by context – by a visible curved pipe or by an occluder whose color indicated the pipe’s shape. This mirrors tracking under occlusion in the real world, where the visual system must often infer trajectory information from context rather than direct observation, as objects frequently move in and out of sight (e.g., a car moving along a road may be temporarily hidden by other vehicles, but its path can be inferred from the road layout). Our experiment was designed to characterize the contribution of such contextual constraints to motion tracking during occlusion. In the pipe condition, trajectory information was always available, but motion characteristics such as speed and direction were not directly perceivable. To isolate the contribution of the pure motion information and dissociate it from the extrapolation of motion, we created two conditions: In the ball-only condition, motion characteristics were directly observable, but trajectory information was only revealed progressively as the ball moved (i.e., contextual motion was unavailable). In the occluder condition, participants had to infer both trajectory and motion characteristics from prior knowledge alone. By cross-decoding across these two conditions, we revealed that contextual information leads to an “ahead of time” coding of occluded motion: Representations of motion direction in the occluder conditions systematically preceded those in the ball-only condition, revealing predictive representations when tracking invisible motion.

The behavioral results converge with the neural findings, as observed with position reports for the ball being predictively biased towards future positions. This effect was particularly pronounced under occlusion, suggesting that when motion characteristics are not directly perceivable, observers have to rely more heavily on prior knowledge for successful tracking. Our results also indicate stronger forward displacement during downward compared to upward motion conditions of the ball. This is consistent with previous results showing that observers tend to report increasing forward velocity for downward motion and decreasing velocity for upward motion (Freyd & Finke, 1984; Hubbard, 2014), consistent with stronger representational momentum in the direction of natural gravity (Freyd & Jones, 1994; Hubbard, 1995). Together, these results suggest that as motion becomes more complex (i.e., requiring predictions about multiple features such as direction and speed), tracking increasingly relies on internalized physical priors. Similar influences of physical priors have been observed in other motion-induced position shifts, such as the flash-lag effect (Hogendoorn, 2020) and frohlich effect (Müsseler et al., 2002).

Together, predictive neural representations and anticipatory behavioral responses could help compensate for sensorimotor delays during active vision and manual action. Although anticipatory coding leads to a greater spatial error in behavioral responses (see Goettker et al., 2021 for a similar result in gaze control), the bias of responses towards future states can counteract sensorimotor delays. Anticipatory coding may thus be beneficial for temporally accurate coding: previous eye-tracking work suggests that observers use contextual information for predictively adjusting gaze when tracking objects in naturalistic situations, allowing them to compensate for sensorimotor delays (Bosco et al., 2015; Goettker et al., 2021; Kowler et al., 2019).

Previous work has shown anticipatory coding in motion tracking as a mechanism to compensate for neural delays when the motion trajectories are linear and one-dimensional (Blom et al., 2020; Hogendoorn & Burkitt, 2018; Turner et al., 2023, 2025). Our findings extend this literature in two respects. First, consistent with prior work, we observe anticipatory representations during the tracking of occluded non-linear motion, suggesting that predictive mechanisms operate even when the moving object is no longer visible. Second, and more critically, these representations persist through later stages of occlusion. This pattern diverges from previous studies, which reported motion extrapolation only during early phases of the occlusion (Teichmann et al., 2022; Yeh et al., 2024). We argue this sustained activity reflects a hallmark of non-linear motion tracking. Unlike linear trajectories, where an initial extrapolation based on starting position and velocity is sufficient to anticipate future states, non-linear trajectories shaped by environmental constraints require ongoing computation: the future position of the object depends not only on its current motion but on how that motion interacts with the context. Sustained neural representations may therefore be necessary to maintain an accurate model of the object’s path as it unfolds along complex, non-linear trajectories. More broadly, the present study examined only one class of environmental constraint - path geometry. However, predictive motion tracking may also incorporate knowledge about physical object properties that influence motion dynamics (Kham et al., 2026). Future work could investigate the range of environmental and object-specific constraints that influences tracking, thereby grounding our understanding of motion tracking in real-world settings.

Our findings are consistent with proposals that alpha rhythms support top-down feedback processes, here reflected in “ahead of time” neural coding during motion tracking based on prior knowledge (Stecher et al., 2025; Turner et al., 2023). An alternative interpretation is that alpha-band activity reflects spatial attention mechanisms (Foster et al., 2017; Samaha et al., 2016). Although we controlled for overt eye movements, covert attentional shifts could, in principle, contribute to the observed tracking effects (Yuasa et al., 2025). However, our data is more consistent with an anticipatory coding account than a purely spatial-attention explanation. In particular, in the case of a strictly position-locked, moment-to-moment updating signal, we should observe an on-diagonal decoding in the temporal generalisation matrices. Instead, we observe a more abstract representation of motion direction as evidenced by the widespread activation that generalizes across conditions. Taken together, our findings suggest that alpha-band dynamics are involved in predictive tracking of motion under occlusion, although they do not fully disentangle anticipatory coding from spatial attention mechanisms. Future work combining spatial cueing paradigms with temporally resolved decoding approaches will be important for more directly dissociating these accounts.

Our findings open several avenues for future research. First, while our stimuli were manipulated along both vertical (up versus down) and horizontal (left versus right) motion axes, we do not observe similar results between them (see Supplementary S3). This raises the question of whether distinct neural mechanisms underlie motion tracking in the vertical axis. One possibility is that alpha dynamics are more strongly engaged during interhemispheric tracking across hemifields than within them (Stephan et al., 2007), suggesting that alpha-band activity may not be a sensitive marker of within-hemifield motion tracking along the vertical axis. Second, our paradigm specifically targets motion direction along a non-linear trajectory. However, motion tracking likely involves other dynamic object properties beyond direction. Future work could examine whether alpha rhythms similarly represent properties such as changes in form, speed, or identity, which would speak to the broader generalizability of alpha rhythms for dynamic visual representation. Third, future work could examine to what extent motion tracking under constraints is driven by prior expectations about the object (e.g., objects with different friction (spiky ball vs smooth ball) will move differently through the same surface). In the present study, participants only needed to infer the behavior of a single falling object under gravity. It remains unclear whether tracking mechanisms would operate similarly when object-specific properties alter expected motion, for instance, causing an object to slow down or stop moving altogether.

Taken together, our findings show that motion tracking for non-linear but predictable trajectories is supported by cortical alpha dynamics. Our behavioral and neural results further provide converging evidence for predictive tracking, with representations anticipating future states of the object. This predictive tracking was particularly pronounced under occlusion, suggesting that the brain more strongly predicts future world states when sensory input is unavailable. Predictive tracking may therefore provide a strategy for retaining temporal precision in a dynamically changing world where objects constantly move in and out of sight.

## Author contributions

Conceptualization: S.A., D.K.. Methodology: S.A., D.K.. Investigation and formal analysis: S.A. Visualization and writing-original draft: S.A. Writing-review and editing: S.A., D.K., F.P., L.-C.Y., K.D, Supervision: D.K., L.-C.Y, K.D. Funding acquisition: D.K., K.D.

## Supplementary material

**S1.**
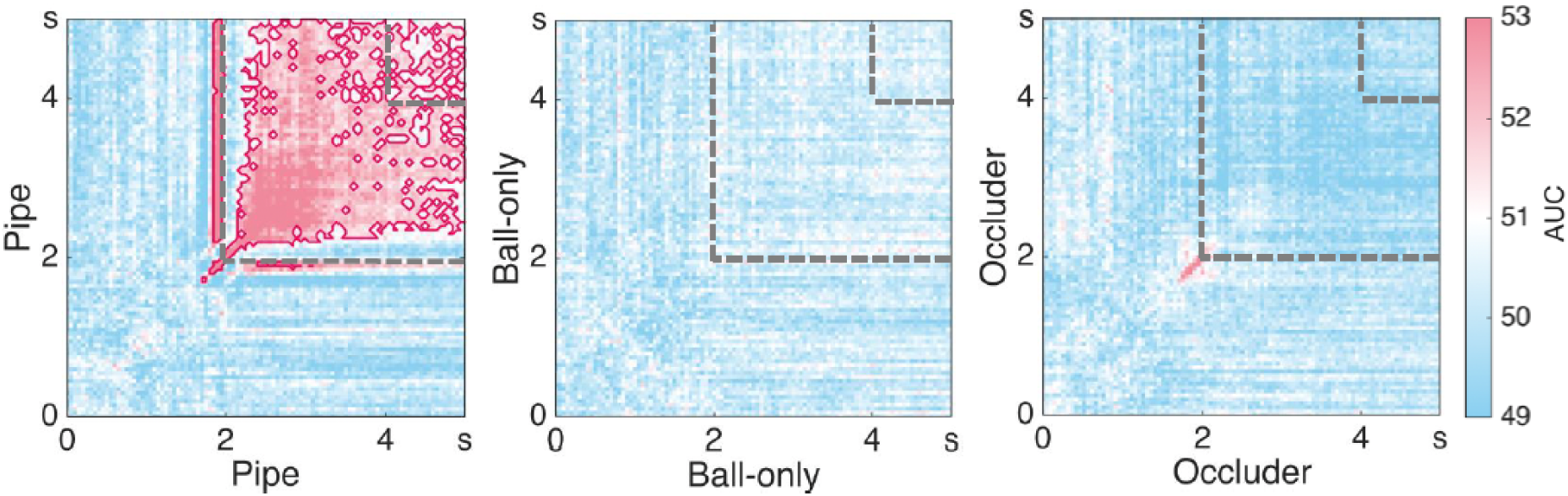
Within condition temporal generalization on evoked potentials (Left vs Right). Linear classifiers were trained and tested on time-locked evoked potentials to decode horizontal motion direction (left versus right). Here, classifiers were trained and tested within the same conditions. Each cell in the matrix reflects decoding accuracy (AUC) at a given train–test time point. The grey dashed lines demarcate the critical time window (2–4 s), spanning the period from which the ball changes direction from its vertical descent until the ball exits. Significant clusters (cluster-based permutation test with TFCE, *p* < 0.05) are outlined in dark pink.

**S2.**
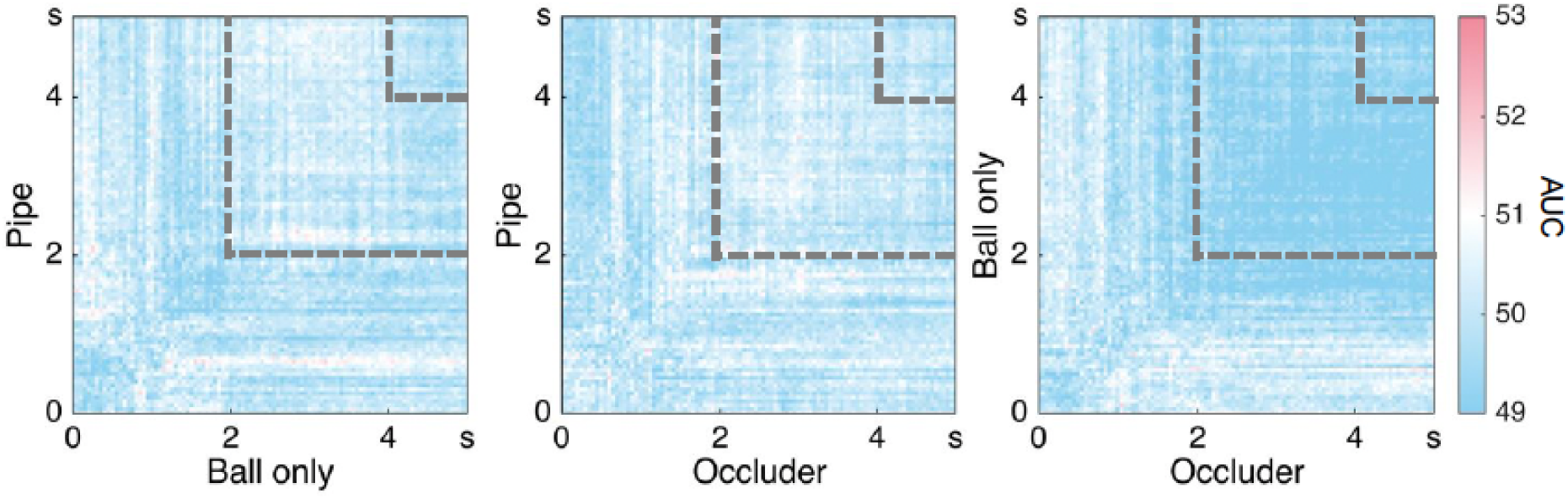
Cross condition temporal generalization on evoked potentials (Left vs Right). Linear classifiers were trained and tested on evoked potentials to decode horizontal motion direction (left versus right). Here the classifier was trained on data from each time point of one condition (e.g., Pipe) and tested it on data from all possible time points of a different condition (e.g., Occluder), resulting in cross-condition temporal generalization matrices. These matrices were symmetrized by averaging across train-test directions (e.g., Train A/Test B and Train B/Test A) for each participant. Individual matrices were then averaged to produce group-level cross-condition generalization matrices. No significant clusters were observed. The grey dashed lines demarcate the critical time window (2–4 s), spanning the period from which the ball changes direction from its vertical descent until the ball exits.

**S3.**
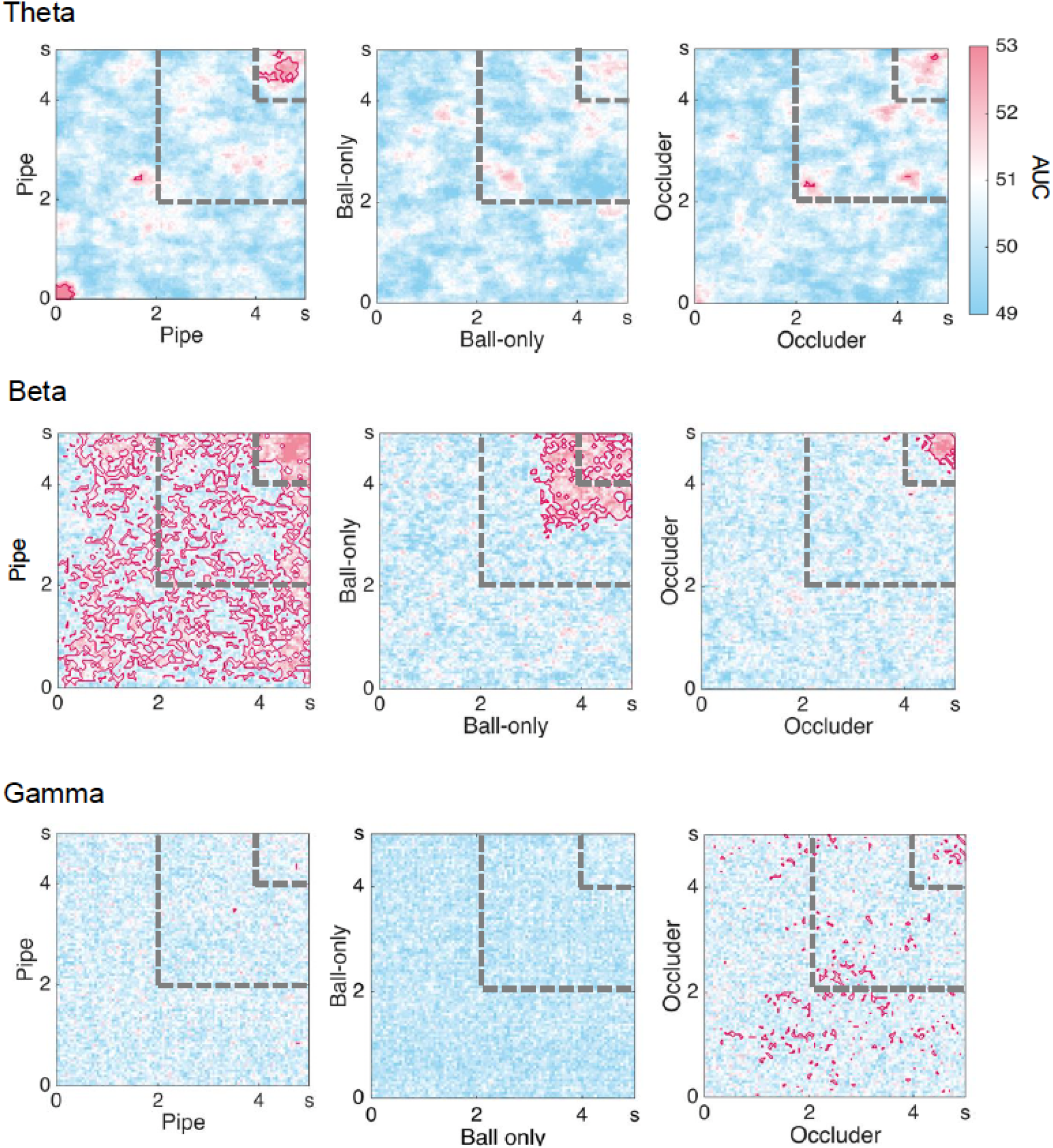
Within-condition temporal generalization in other frequency bands (Left vs Right). Linear classifiers were trained and tested on alpha power (8–12 Hz) patterns to decode horizontal motion direction (left versus right). Here, classifiers were trained and tested within the same conditions. Each cell in the matrix reflects decoding accuracy (AUC) at a given train–test time point. The grey dashed lines demarcate the critical time window (2–4 s), spanning the period from which the ball changes direction from its vertical descent until the ball exits. Significant clusters (cluster-based permutation test with TFCE, *p* < 0.05) are outlined in dark pink.

**S4.**
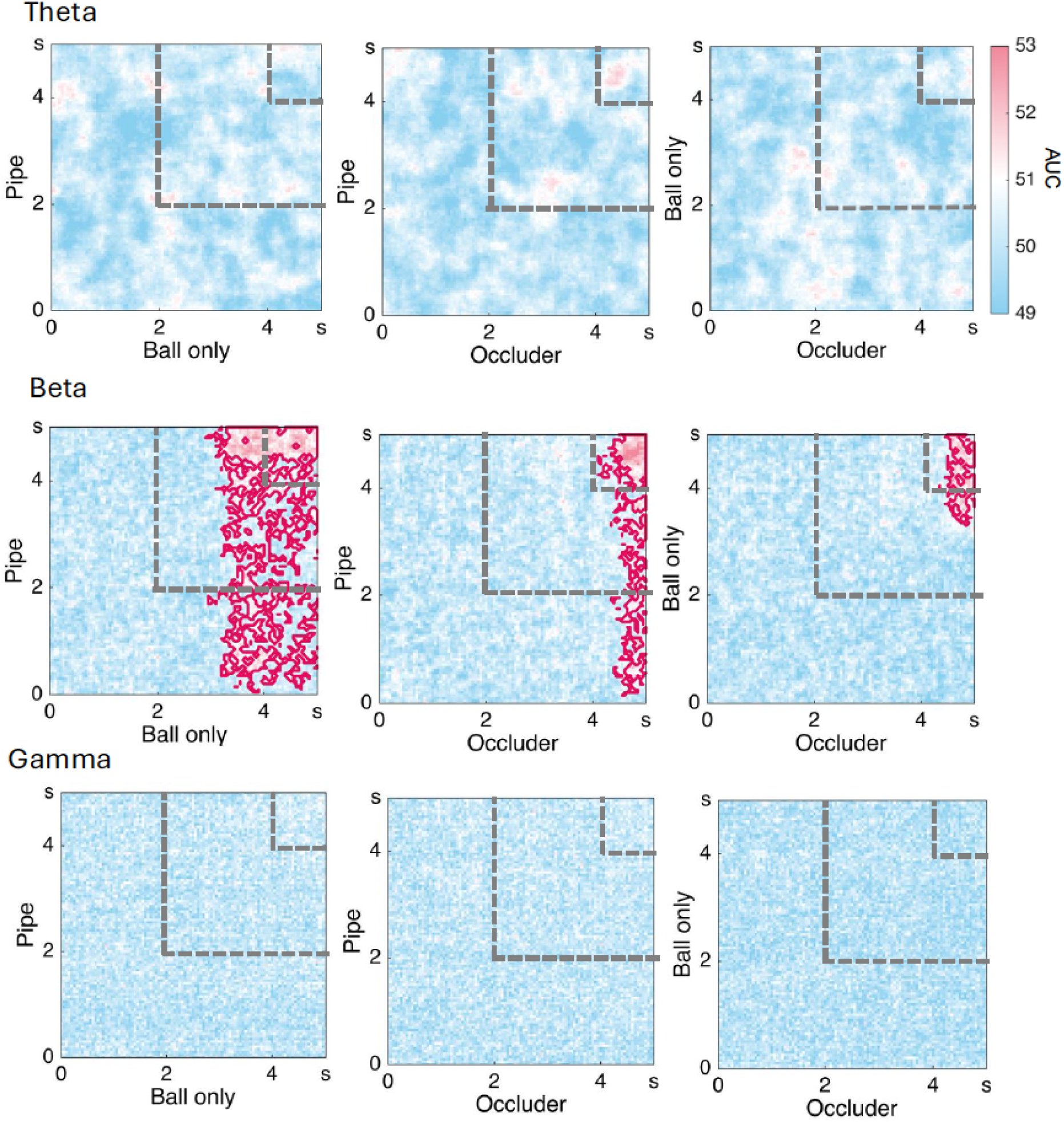
Cross-condition temporal generalization in other frequency bands (Beta, Theta, Gamma) (Left vs Right). Linear classifiers were each trained and tested on the power patterns of different frequency bands (beta: 4-7Hz; theta: 13-30Hz; gamma: 31-70Hz) to decode horizontal motion direction (left versus right). Here, the classifier was trained on data from each time point of one condition (e.g., Pipe) and tested it on data from all possible time points of a different condition (e.g., Occluder), resulting in cross-condition temporal generalization matrices. The matrices were averaged across train–test directions for each participant and then averaged across participants. Each cell in the matrix reflects decoding accuracy (AUC) at a given train–test time point. The grey dashed lines demarcate the critical time window (2–4 s), spanning the period from which the ball changes direction from its vertical descent until the ball exits. Significant clusters (cluster-based permutation test with TFCE, *p* < 0.05) are outlined in dark pink. No significant shared clusters were observed within the time period of interest. Note that any significant decoding beyond 4s likely reflects a sensory-evoked response to the ball’s reappearance. Additionally, decoding in the Pipe condition is strongly driven by the presence of the pipe shape throughout the trial.

**S5.**
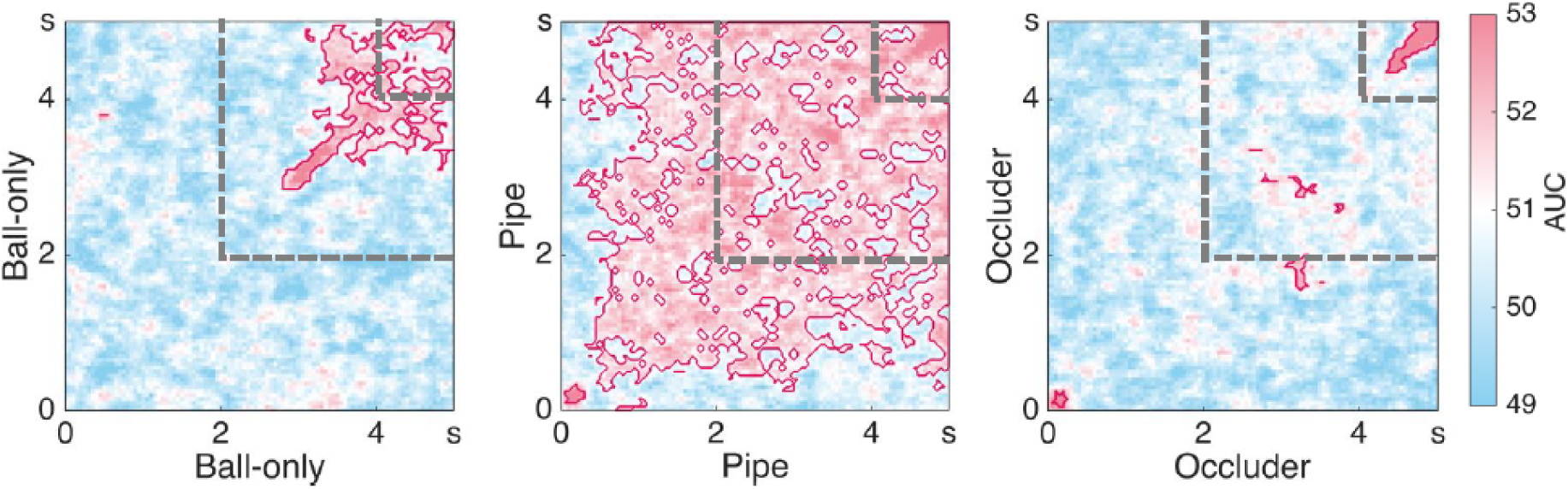
Within-condition temporal generalization (Alpha band) (Up vs Down). Linear classifiers were trained and tested on alpha power (8–12 Hz) patterns to decode vertical motion direction (up versus down). Here, classifiers were trained and tested within the same conditions. Each cell in the matrix reflects decoding accuracy (AUC) at a given train–test time point. The grey dashed lines demarcate the critical time window (2–4 s), spanning the period from which the ball changes direction from its vertical descent until the ball exits. Significant clusters (cluster-based permutation test with TFCE, *p* < 0.05) are outlined in dark pink.

**S6.**
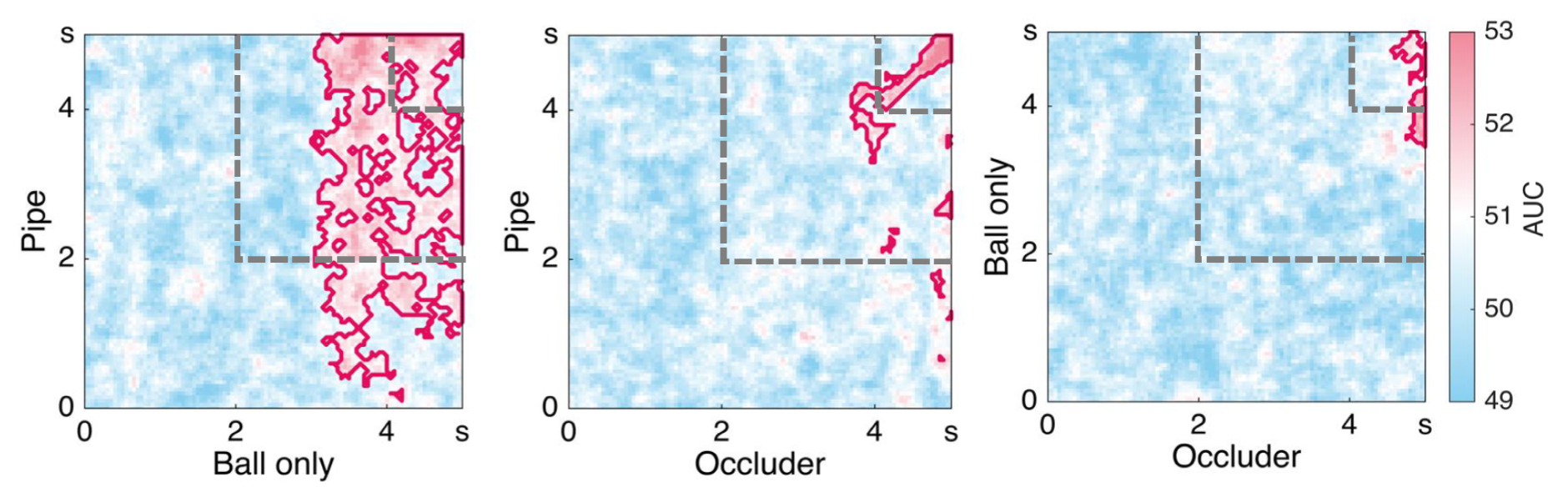
Cross-condition temporal generalization (Alpha band) (Up vs Down). Linear classifiers were trained and tested on alpha power (8–12 Hz) patterns to decode vertical motion direction (up versus down). Here, classifiers were trained on data from each time point of one condition (e.g., Pipe) and tested on data from all possible time points of a different condition (e.g., Occluder), resulting in cross-condition temporal generalization matrices. The matrices were averaged across train–test directions for each participant and then averaged across participants. Each cell in the matrix reflects decoding accuracy (AUC) at a given train–test time point. The grey dashed lines demarcate the critical time window (2–4 s), spanning the period from which the ball changes direction from its vertical descent until the ball exits. Significant clusters (cluster-based permutation test with TFCE, *p* < 0.05) are outlined in dark pink. No significant shared clusters emerged during the critical window across conditions.

**S7.**
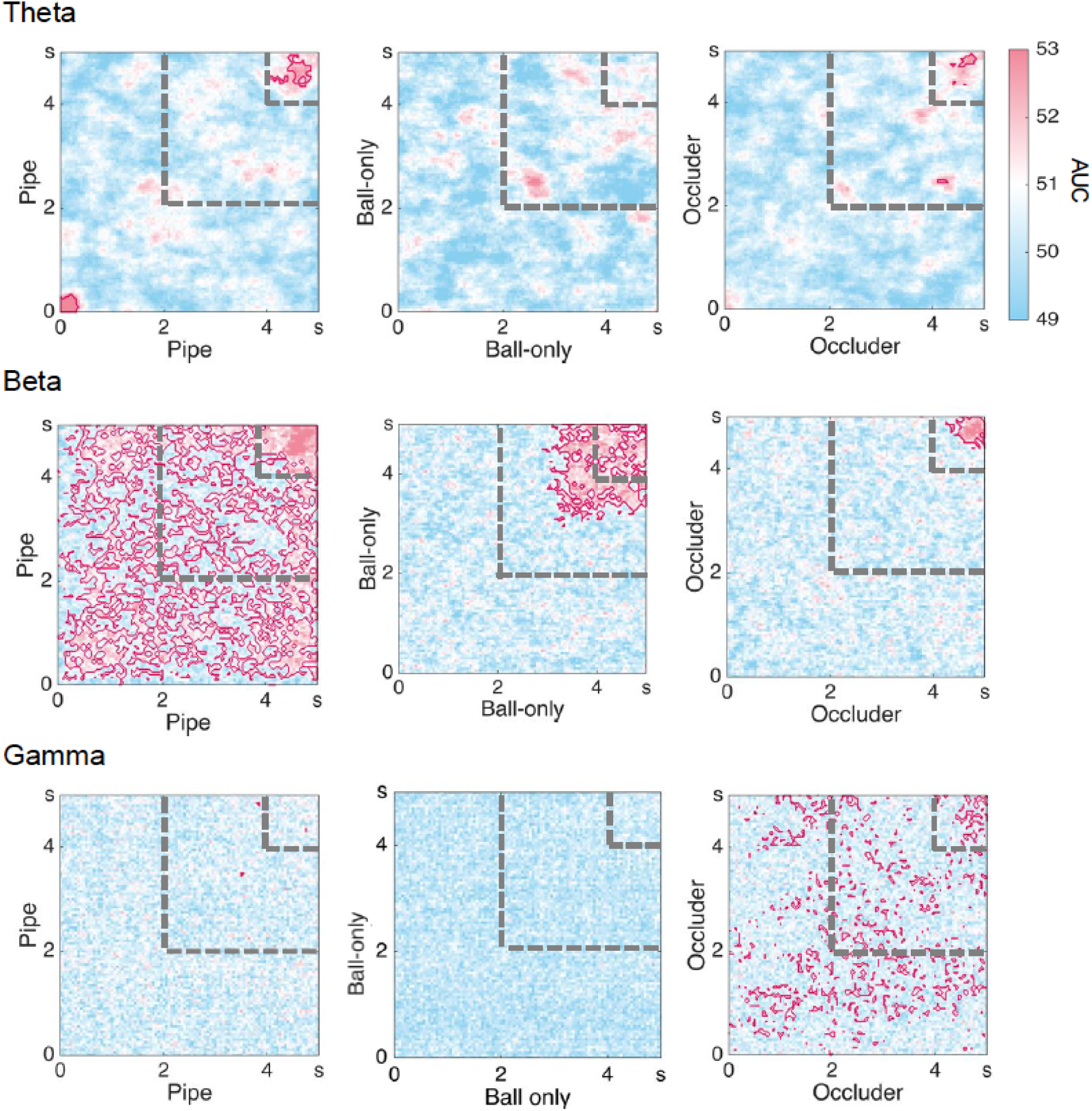
Within-condition temporal generalization in other frequency bands (Theta, Beta, Gamma) (Up vs Down). Linear classifiers were each trained and tested on the power patterns of different frequency bands (beta: 4-7Hz; theta: 13-30Hz; gamma: 31-70Hz) to decode vertical motion direction (up versus down). Here, classifiers were trained on data from each time point of one condition (e.g., Pipe) and tested on data from all possible time points of a different condition (e.g., Occluder), resulting in cross-condition temporal generalization matrices. The matrices were averaged across train–test directions for each participant and then averaged across participants. Each cell in the matrix reflects decoding accuracy (AUC) at a given train–test time point. The grey dashed lines demarcate the critical time window (2–4 s), spanning the period from which the ball changes direction from its vertical descent until the ball exits. Significant clusters (cluster-based permutation test with TFCE, *p* < 0.05) are outlined in dark pink. No significant shared clusters emerged during the critical window across conditions.

